# Early life maturation of human visual system white matter is altered by monocular enucleation

**DOI:** 10.1101/690701

**Authors:** Benjamin T. Dunkley, Marlee Vandewouw, Arijit Chakraborty, Margot J. Taylor, Brenda Gallie, Daphne L. McCulloch, Benjamin Thompson

**Author notes:** Co-first authors (Benjamin T. Dunkley, Marlee Vandewouw). **Corresponding author:** Benjamin T. Dunkley, PhD, Department of Diagnostic Imaging, Hospital for Sick Children, 555 University Ave. Toronto, M5G 1X8, Ontario, Canada, ex 309117.

## Abstract

Monocular enucleation early in life and the resultant lack of binocular visual input during visual development results in functional and structural brain changes in adulthood, including alterations in white matter microstructure. However, the time courses of these neurodevelopmental changes are unknown. Here, we investigated whether structural brain changes were present at 8 to 12 years of age in a group of children with a history of monocular enucleation prior to 3 years of age (the ME group) relative to control participants with normal binocular vision (the BC group). Structural connectivity was measured using diffusion tensor imaging (DTI). Relative to the BC group, the ME group exhibited significantly increased radial and mean diffusivity in the optic radiation contralateral to the enucleated eye, the bidirectional interhemispheric V1 to V1 tracts and the V1 to MT tract ipsilateral to the enucleated eye. These changes indicate abnormal myelinization and reduced axonal density in subcortical and cortical visual pathway white matter structures following unilateral enucleation and loss of binocular vision. Our findings are broadly consistent with those recently reported for older uniocular individuals suggesting that these effects are present in childhood and persist into adulthood.

## Introduction

Binocular vision plays an important role in guiding visual cortex development. In juvenile, non-human animals, experimental manipulations of binocular vision involving monocular eye-lid suture, induction of anisometropia (unequal refractive error between the two eyes) or strabismus (eye misalignment), cause widespread neuroplastic changes within the primary and extrastriate visual cortex (Mitchell & Sengpiel, 2018). Humans may also experience binocular vision disruption during infancy due to naturally occurring anisometropia, strabismus, or cataract. These pediatric eye conditions often result in amblyopia, a neurodevelopmental disorder of vision associated with monocular and binocular visual dysfunction (Holmes & Clarke, 2006, Maurer & Mc, 2018). In agreement with the non-human animal literature, a range of neuroanatomical changes have been reported within the visual pathway of humans with amblyopia. These include disrupted white matter structure (Allen, Schmitt, Kushner & Rokers, 2018, Allen, Spiegel, Thompson, Pestilli & Rokers, 2015, Duan, Norcia, Yeatman & Mezer, 2015, Li, Jiang, Guo, Li, Cai & Yin, 2013, Qi, Mu, Cui, Li, Shi, Liu, Xu, Zhang, Yang & Yin, 2016, Xie, Gong, Xiao, Ye, Liu, Gan, Jiang & Jiang, 2007, Zhai, Chen, Liu, Zhao, Zhang, Luo & Gao, 2013), reduced subcortical and cortical grey matter density (Mendola, Conner, Roy, Chan, Schwartz, Odom & Kwong, 2005, Xiao, Xie, Ye, Liu, Gan, Gong & Jiang, 2007), and alterations in effective (Li, Mullen, Thompson & Hess, 2011) and resting state connectivity (Ding, Liu, Yan, Lin & Jiang, 2013, Liang, Xie, Yang, Yin, Wang, Yu, He & Wang, 2017, Mendola, Lam, Rosenstein, Lewis & Shmuel, 2018, Wang, Li, Guo, Peng, Li, Qin & Yu, 2014). However, it is unclear whether these changes are due to lost binocular visual input *per se* or because of mismatched, competing inputs from the two eyes during visual development.

Individuals who have experienced early-life monocular enucleation, typically as a treatment for retinoblastoma, represent an alternative way to study the impact of binocular vision loss on visual system development (Steeves, Gonzalez & Steinbach, 2008). Because the input from one eye has been lost entirely, any changes in visual function or neuroanatomy must be due to a loss of visual function *per se*. There is a substantial literature on the effects of early-life monocular enucleation on visual function (Kelly, Moro & Steeves, 2012, Steeves et al., 2008). Uniocular individuals have largely normal monocular vision and those enucleated early (before 4 years of age) showed improved contrast sensitivity compared to binocular or uniocular viewing by typical binocular controls (Nicholas, Heywood & Cowey, 1996). However, aspects of motion and face perception that rely on higher-level, binocular regions of the extrastriate visual cortex, are impaired (Gonzalez, Lillakas, Greenwald, Gallie & Steinbach, 2014, Kelly, Gallie & Steeves, 2012, Kelly, Zohar, Gallie & Steeves, 2013, Steeves, Gonzalez, Gallie & Steinbach, 2002). Interestingly, changes in multimodal sensory processing involving improved auditory direction discrimination and a relative shift away from vision in auditory/visual integration tasks have also recently been reported in adults with a history of early-life monocular enucleation (Hoover, Harris & Steeves, 2012, Moro & Steeves, 2012, Moro & Steeves, 2013). These results suggest multimodal compensation for early uniocular vision loss.

Recently, a number of studies have investigated brain structure and function in adults with a history of early-life monocular enucleation using magnetic resonance imaging. These studies have revealed the following pattern of brain structure alterations: reduced optic tract and LGN volumes contralateral to the enucleated eye (Kelly, McKetton, Schneider, Gallie & Steeves, 2014), larger surface area and a more complex sulcal structure in V1 ipsilateral to the enucleated eye along with larger auditory and multisensory areas in both hemispheres (Kelly, DeSimone, Gallie & Steeves, 2015), an asymmetry in the medial geniculate body (MGB; involved in auditory processing) whereby the left MGB is larger than the right MGB (Moro, Kelly, McKetton, Gallie & Steeves, 2015), and, reduced functional activation of extrastriate areas involved in face perception (Kelly, Gallie & Steeves, 2018). Therefore, a complete loss of binocular vision early in life appears to have effects on the structure and function of brain areas involved in vision, audition and multisensory integration. This is consistent with the widespread effects of early-life monocular enucleation on brain development in non-human animal models (Nys, Scheyltjens & Arckens, 2015).

Diffusion tensor imaging has also been used to examine white matter structure in adults with a history of uniocular enucleation. DTI uses MRI pulse sequences that exploit the diffusion of water molecules along relatively unconstrained spatial gradients to map structural connections in the brain (Jellison, Field, Medow, Lazar, Salamat & Alexander, 2004). Principally, it can resolve white matter tractography to give a ‘circuit diagram’ of the central nervous system. Briefly, water will diffuse more easily along an axon than in radial directions – this motion can be modelled as a tensor that describes the 3-dimensional movement of the molecules. Properties of diffusion characteristics can be extracted from tensor models to provide information regarding the underlying white matter microstructure within the brain (Alexander, Hurley, Samsonov, Adluru, Hosseinbor, Mossahebi, Tromp do, Zakszewski & Field, 2011). Common parameters include: axial diffusivity (AD) which is related to the vector length of the tensor, or the amount of diffusion in the primary direction – this increases with maturation and decreases with axonal injury; radial diffusivity (RD), a measure of diffusion orthogonal to the primary direction, which decreases with maturation and increases with demyelination or white matter injury; mean diffusivity (MD), the total diffusion of water content in a voxel, which decreases with maturation; and fractional anisotropy (FA), a measure of the diffusion constraint in a voxel. FA increases with maturation and is proportional to axonal density and diameter.

DTI measures in adults with a history of early-life monocular enucleation have revealed a range of white matter alterations in the visual cortex, some of which are evident only as changes in the lateralization of white matter parameters relative to those of binocular controls (Wong, Rafique, Kelly, Moro, Gallie & Steeves, 2018). Specifically, uniocular adults exhibited higher FA values in the optic radiations ipsilateral to the enucleated eye, whereas controls had higher FA contralateral to the non-dominant eye (the non-dominant eye was referenced to the enucleated eye in patients). Similarly, in the V1 to LGN projections, uniocular adults had higher RD in the tracts contralateral to the enucleated eye, whereas controls had higher RD ipsilateral to the non-dominant eye. There was also one statistically significant between-group difference; uniocular adults had lower FA than controls within their interhemispheric V1 to V1 projections.

The vast majority of brain imaging work in uniocular participants has been conducted with adults. One previous study measured visual cortex white matter structure in children with retinoblastoma, some of whom had a history of monocular enucleation (Barb, Rodriguez-Galindo, Wilson, Phillips, Zou, Scoggins, Li, Qaddoumi, Helton, Bikhazi, Haik & Ogg, 2011). The age-related development of white matter appeared to follow a normal trajectory in children with retinoblastoma, however the study was not designed to enable a detailed analysis of white matter microstructure changes following monocular enucleation.

Building on this prior work, the aim of this study was to assess whether changes in visual pathway white matter structure associated with early-life monocular enucleation are evident in late childhood when a range of higher-level visual functions are still developing. We used DTI to assess both feedforward and feedback fibers within the optic radiations connecting the lateral geniculate nucleus and V1, the interhemispheric V1 to V1 tracts and the within-hemisphere V1 to MT tracts. The latter tracts were of interest because of previously reported motion perception abnormalities in uniocular adults (Gonzalez et al., 2014, Kelly et al., 2013, Steeves et al., 2002).

## Methods

### Materials and Methods

#### Participants

Six children with a history of monocular enucleation for the treatment of retinoblastoma (the ME group, age range 8-13 years) and six age-matched binocular controls (the BC group, age range 9-12 years) participated. All participants had normal or corrected-to-normal visual acuity. After receiving a complete description of the study, all participants gave informed written consent. Imaging was performed at the Hospital for Sick Children, after approval from the institutional Research Ethics Board. For both groups, exclusion criteria included: a history of any neurological disorder; presence of intra-corporeal metal and/or medical devices contraindicated for MRI recording.

#### Procedures

##### Imaging protocol

All imaging was performed using a Siemens 3.0T MAGNETOM PrismaFIT scanner and 20 channel head and neck coil. T1-weighted images were collected using a 3D magnetization prepared rapid gradient echo (MPRAGE) pulse sequence (TR/TE: 2300/2.98ms; FA: 9°; FOV: 192 × 240 × 256 mm; resolution: 1.0 mm^3^ isotropic; scan time: 5 min). Multi-shell diffusion images were collected using an echo planar imaging (EPI) diffusion pulse sequence TR/TE: 3800/73 ms; FA: 90°; FOV: 244 × 244 × 140 mm; 2.0 mm isotropic voxels; b = 1000/1600/2600 s/mm2 (30/40/60 directions); 15 interleaved b = 0 s/mm2 volumes; scan time: 10.25 min).

##### Data preprocessing

###### DTI

The T1-w anatomical images were skull-stripped using the Freesurfer image analysis suite, which is documented and freely available for download (http://surfer.nmr.mgh.harvard.edu/). FSL tools (Jenkinson, Bannister, Brady & Smith, 2002) and custom scripts were used to correct for eddy current distortions and head motion. The mean for each b-value (0, 1000, 1600, 2600) was calculated, and transformations between each mean b-value image and the mean b = 0 image were obtained using FSL’s FLIRT (Jenkinson et al., 2002). Each individual volume was first registered to the mean of its b-value using FSL’s FLIRT (Jenkinson et al., 2002), and then to the mean b = 0 image using the transformations, while adjusting the corresponding b-vectors. A transformation between MNI space and diffusion space was obtained via FSL’s FNIRT by registering the MNI template to the mean b = 0 image. FSL’s FDT was used to fit the diffusion tensors, obtaining fraction anisotropy (FA), radial diffusivity (RD), axial diffusivity (AD), and mean diffusivity (MD) images. FA, RD, AD, and MD images were flipped as necessary so that the remaining eye was on the right in the uniocular group.

###### TBSS analysis

Voxelwise statistical analysis of the FA, RD, AD, and MD images was performed using tract-based spatial statistics, TBSS (Smith, Johansen-Berg, Jenkinson, Rueckert, Nichols, Miller, Robson, Jones, Klein, Bartsch & Behrens, 2007). A study-specific FA template was generated following nonlinear co-registration. The template was skeletonized, thresholded at 0.2, and projected onto each subject’s FA, RD, AD, and MD images.

###### Probabilistic tractography

FSL’s BEDPOSTX (Jbabdi, Sotiropoulos, Savio, Grana & Behrens, 2012) was used to estimate the diffusion parameters using the multi-shell model. Probabilistic tractography using FSL’s PROBTRACKX (Behrens, Berg, Jbabdi, Rushworth & Woolrich, 2007, Behrens, Woolrich, Jenkinson, Johansen-Berg, Nunes, Clare, Matthews, Brady & Smith, 2003) was employed to reconstruct the feedforward LGN to V1 projection (LGN source -> V1 termination), projection of V1 source -> LGN termination, interhemispheric V1 projections, and V1 source -> MT+ termination in each hemisphere. Bilateral masks of the primary visual cortex (V1), middle temporal visual area (MT+), lateral geniculate body (LGN), and cerebral white matter (WM) were extracted from the Juelich Histological atlas (Eickhoff, Heim, Zilles & Amunts, 2006, Eickhoff, Paus, Caspers, Grosbras, Evans, Zilles & Amunts, 2007, Eickhoff, Stephan, Mohlberg, Grefkes, Fink, Amunts & Zilles, 2005) and the Harvard-Oxford subcortical structural atlas (Desikan, Segonne, Fischl, Quinn, Dickerson, Blacker, Buckner, Dale, Maguire, Hyman, Albert & Killiany, 2006, Frazier, Chiu, Breeze, Makris, Lange, Kennedy, Herbert, Bent, Koneru, Dieterich, Hodge, Rauch, Grant, Cohen, Seidman, Caviness & Biederman, 2005) in MNI space, and each subject’s MNI to diffusion non-linear transformations were inputted. Default settings were used, with the addition of distance correction and modified Euler streamlining to improve accuracy. Termination masks were also selected as waypoint masks, and where applicable, WM was used as an exclusion mask. The resulting eight tracts (four per hemisphere) were masked by the TBSS skeleton, and FSL tools (Jenkinson, Beckmann, Behrens, Woolrich & Smith, 2012) were used to extract the average FA, RD, AD, and MD values.

###### Statistical analysis

For TBSS, group differences between the binocular (BC) and monocular enucleation (ME) subjects were analysed using FSL’s randomise function (Winkler, Ridgway, Webster, Smith & Nichols, 2014), using 5000 permutations and threshold-free cluster enhancement. TBSS’s symmetry tool was also used to test group differences in hemispheric asymmetries in diffusion characteristics.

Between group differences in FA, RD, AD, and MD for tracts ipsilateral and contralateral to the enucleated eye were tested using the non-parametric Mann-Whitney U test. Additionally, differences between tracts contralateral vs. ipsilateral to the enucleated eye were tested for the ME group using Wilcoxen signed ranks tests. We did not perform within group comparisons for the BC group because eye dominance may influence the direction of interhemispheric differences in white matter structure (Wong et al., 2018) and eye dominance data were not collected. The False Discovery Rate (FDR) correction was applied to p values *within* each DTI metric *across* tracts to control for multiple comparisons, and significance was held at *q* < 0.05 (corrected *p*).

## Results

### Patient characteristics

Patients had a mean age of enucleation of 21 months (1.8 years) and an average age of 9.8 years at the time of scanning (**Table 1**). The binocular control group had a mean age of 11 years (9-14 years).

**Table 1.** Characteristics of the children with a history of monocular enucleation.

| Participants | Age at enucleation<br>months (years) | Age at scan<br>(years) | Enucleated eye | Visual acuity<br>(logMAR) |
| --- | --- | --- | --- | --- |
| ME1 | 19.1 (1.59) | 9.58 | Right | -0.01 |
| ME2 | 17.0 (1.42) | 12.42 | Right | 0.06 |
| ME3 | 26.3 (2.91) | 10.03 | Right | -0.14 |
| ME4 | 36.4 (3.03) | 9.56 | Left | -0.14 |
| ME5 | 7.0 (0.58) | 8.46 | Left | -0.28 |
| ME6 | 13.0 (1.08) | 8.83 | Left | -0.3 |
| **Mean** | **21.2 (1.77)** | **9.81** |  | **-0.14** |

### Anatomy of the pathways

We identified the four pathways of interest, LGN-V1 and V1-LGN (the optic radiations), V1-V1 and V1-MT+, in all participants using probabilistic tractography. **Figure 1** shows pathway anatomy for 2 representative participants, one from the BC group (top row), and the ME group (bottom row).

**Figure 1.**
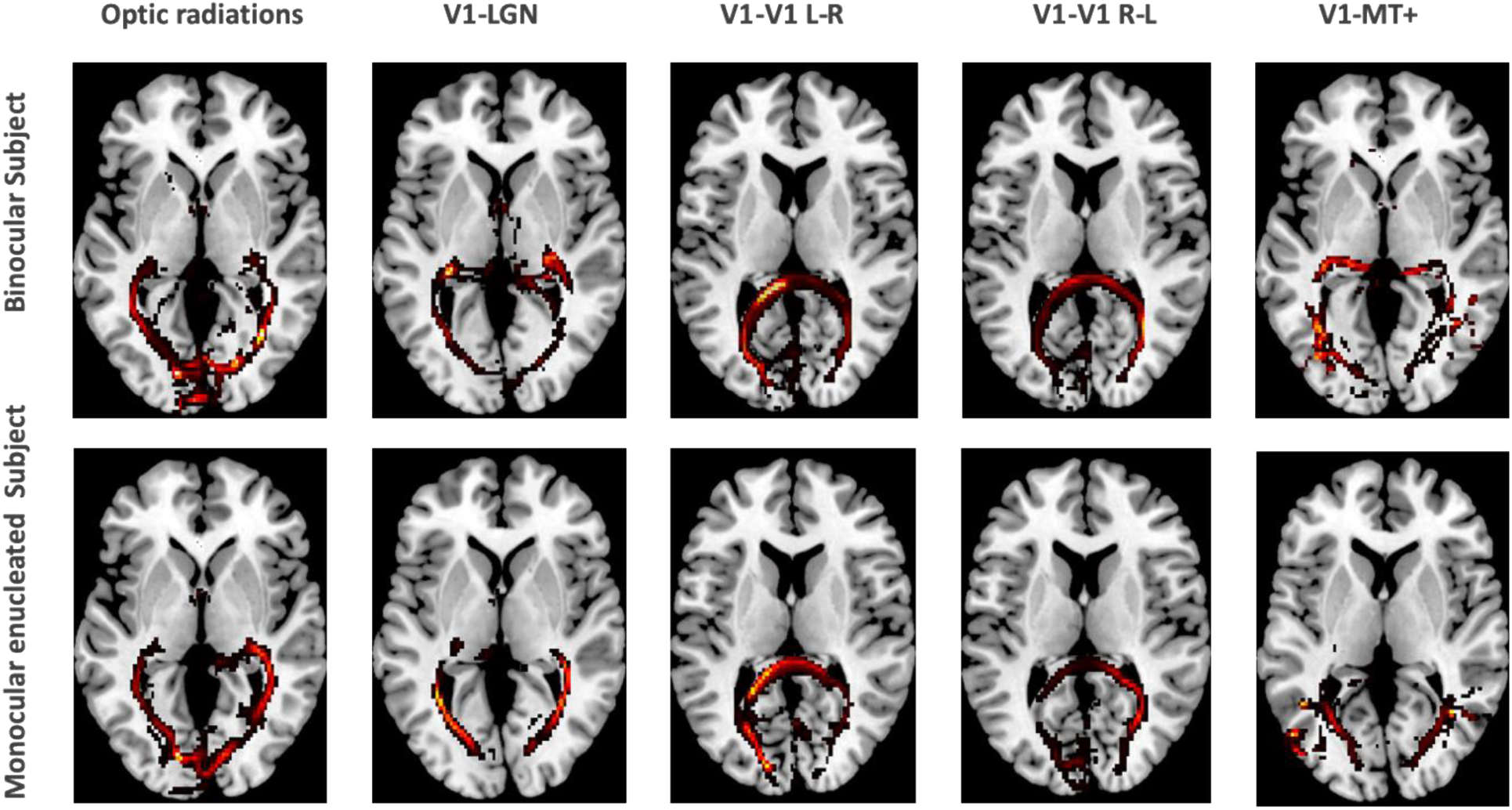
White matter pathways reconstructions calculated using PROBTRACKX of two participants, one from the BC group and one from the ME group, showing LGN-V1 (optic radiations), V1-LGN, interhemispheric V1 (seed in left hemisphere = L-R; seed in right hemisphere = R-L), and V1-MT+ connections, overlaid on a template brain.

### Between group differences

There were no significant differences between groups for TBSS measures of FA, RD, AD or MD. Three significant between group differences were observed when individual tracts identified using probabilistic tractography were analyzed (**Figure 2**). The ME group exhibited significantly increased RD and MD relative to the BC group in the V1-LGN projections contralateral to the enucleated eye (RD: U = 2.5, p = 0.009; MD: U = 2, p = 0.009), the interhemispheric V1 to V1 projections (both contralateral to ipsilateral and ipsilateral to contralateral; RD: U = 3, p = 0.015; MD: U = 3, p = 0.015) and the intrahemispheric V1 to V5 projections ipsilateral to the enucleated eye (RD: U = 1.5, p = 0.004; MD: U = 1.5, p = 0.004). Increased RD for the V1 to V5 projection contralateral to the enucleated eye was also observed, but this difference did not survive FDR correction. Significant between group differences in RD and MD were accompanied by an appropriate trend for reduced FA in the ME group (**Figure 2**), however, these differences did not survive FDR correction.

**Figure 2.**
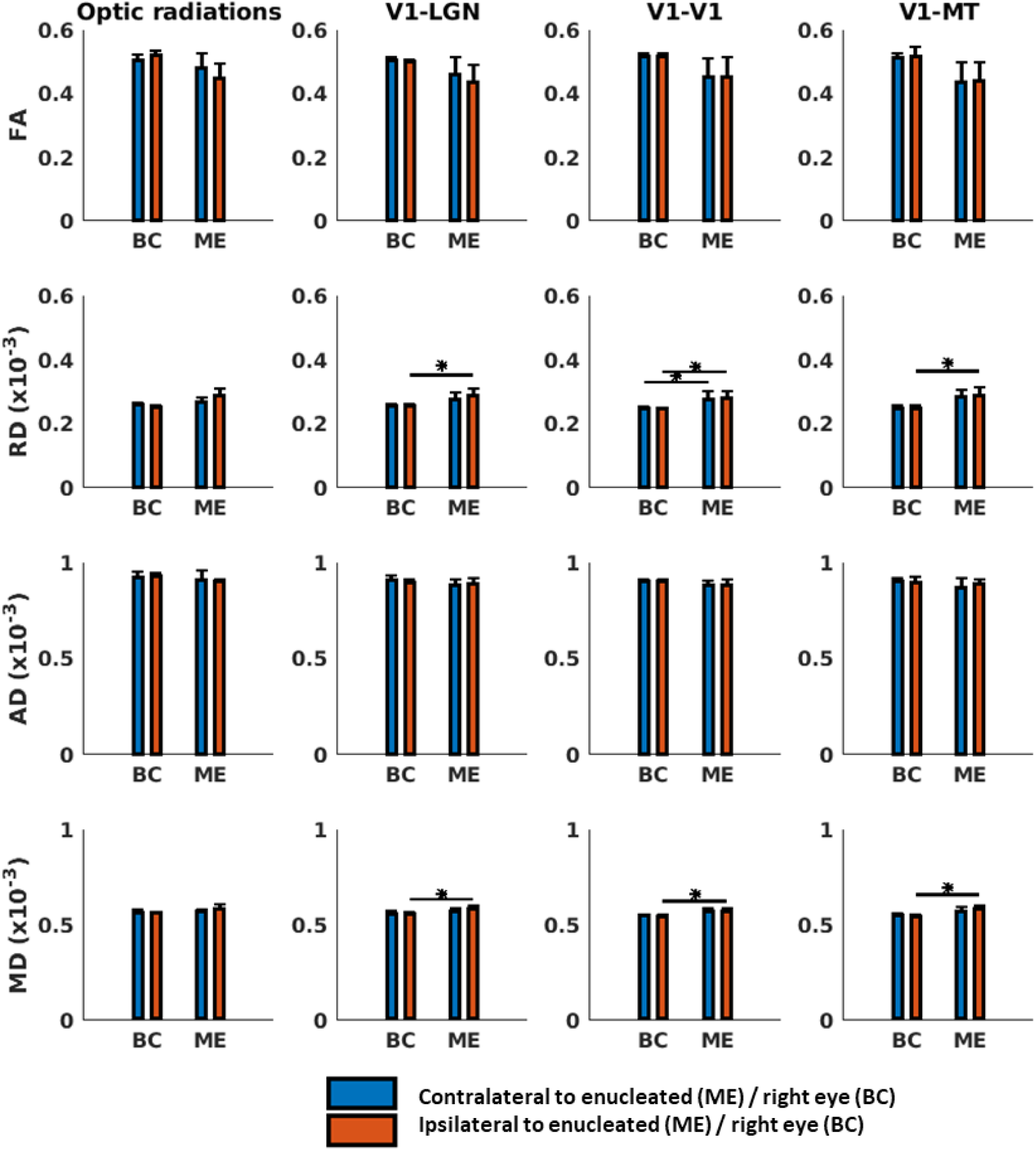
Comparison of DTI measures for visual tracts, including (A) LGN to V1, (B) V1-LGN, (C) interhemispheric V1, and (D) intrahemispheric V1-MT. Tracts were identified as contralateral (blue bars) vs. ipsilateral (orange bars) to the enucleated eye for the ME group and contralateral vs. ipsilateral to the right eye for controls. Error bars depict ±1 SE. * = p < 0.05, FDR corrected.

### Within group differences

No significant differences between tracts contralateral vs. ipsilateral to the remaining eye were observed for the ME group.

## Discussion

In agreement with recent work in adults (Wong et al., 2018), we observed subtle, localized, differences in white matter structure between our ME and BC groups. These differences highlight specific regions of the visual pathway that require binocular input for normal development. All of the statistically significant effects that we observed involved increased RD and MD in the children with early enucleation relative to the typically developing children. Increased RD occurs in the presence of reduced or abnormal myelinization that can impair neural conductivity (Alexander et al., 2011, Song, Sun, Ramsbottom, Chang, Russell & Cross, 2002, Wu, Field, Duncan, Samsonov, Kondo, Tudorascu & Alexander, 2011). Myelinization is directly influenced by neural activity within white matter tracts (Mount & Monje, 2017). Therefore, tracts in the ME group that had increased RD relative to controls may be particularly sensitive to the significant reduction in neural drive that occurs when one eye is removed. Increased MD is associated with reduced axonal density (Alexander et al., 2011), suggesting that both myelinization and tract morphology are affected by uniocular vision and loss of input to the binocular system.

We observed RD and MD increases for both subcortical and cortical tracts. The subcortical changes occurred for feedback projections from V1 to the LGN ipsilateral to the remaining eye in the ME group. Feedforward LGN to V1 projections exhibited a similar trend that did not reach statistical significance. The optic radiations (the white matter tracts that connect the LGN and V1) contralateral to the enucleated eye have the largest reduction in innervation following monocular enucleation because there is a proportionally greater amount of crossed relative to uncrossed ganglion cell axons in each eye (see Kelly et al., 2014 for a discussion). Crossed ganglion cell axons project to the contralateral hemisphere. Furthermore, adults with a history of early life monocular enucleation exhibit a large asymmetry in LGN volume whereby the LGN contralateral to the enucleated eye is smaller (Kelly et al., 2014). Postnatal changes in neural connectivity within the LGN and/or feedback from V1 to the LGN may explain this effect (Kelly et al., 2014). Our observations are in agreement with these previous findings, indicating that subcortical changes associated with early monocular enucleation are present in childhood.

Regarding cortical tracts, the ME group exhibited significantly increased RD and MD in bidirectional interhemispheric V1 to V1 fibers compared to the BC group. This is consistent with Wong et al. who reported significantly reduced FA in bidirectional V1 to V1 fibers in their adult uniocular group. Reduced FA is consistent with increased RD and MD, although FA is a more general measure of white matter microstructure (Alexander et al., 2011). Intracortical connectivity within the normal visual pathway facilitates binocular vision, enables integrated processing across the left and right visual hemifields (Peiker, Wunderle, Eriksson, Schmidt & Schmidt, 2013, Restani & Caleo, 2016, Song & Rees, 2018) and facilitates interhemispheric synchronization that supports sparse encoding of visual information (Knyazeva, 2013). Therefore down-regulation of interhemispheric tracts could affect a variety of visual functions including those known to be disrupted by early life enucleation such as face perception and motion integration. Our results are consistent with a large literature involving non-human animals demonstrating that a range of manipulations including early monocular enucleation or eyelid suture alter the development of intracortical primary visual cortex connections (Nys et al., 2015, Restani & Caleo, 2016).

Increased RD and MD was also evident within the V1 to V5 tract ipsilateral to the enucleated eye with a similar, but not statistically significant trend evident in the contralateral tract. V5 is primarily binocular and therefore may be particularly susceptible to a loss of binocular vision early in life (Born & Bradley, 2005, El-Shamayleh, Kiorpes, Kohn & Movshon, 2010, Ho, Giaschi, Boden, Dougherty, Cline & Lyons, 2005, Thompson, Villeneuve, Casanova & Hess, 2012) and the resulting downregulation of neural input. In addition, V1 to V5 connections are essential for the processing of integrated, global motion (Ajina, Kennard, Rees & Bridge, 2015). Therefore, alterations in these tracts may underpin the motion perception deficits that have been identified in adults with a history of early monocular enucleation. No previous studies have investigated V1 to V5 projections in cases of early monocular enucleation.

Unlike Wong et al. (2015), we did not observe any significant asymmetries within the ME group for ipsilateral vs. contralateral to the remaining eye in any DTI parameters. This cannot be explained by differences between the two studies in the age at which enucleation was performed (mean of 1.7 years for this study and 2 years for Wong et al.). It is conceivable that these asymmetries develop with increasing age to compensate for asymmetrical input to the two hemispheres from the remaining eye.

Overall, our results in children with a history of early monocular enucleation are broadly consistent with those of Wong et al. (2015) in adults; subcortical white matter microstructure is abnormal contralateral to the enucleated eye and intracortical V1 to V1 projections are compromised. Our results demonstrate that these changes are present in childhood and, based on Wong et al’s (2015) data, persist into adulthood. Although white matter changes within the visual pathway are not widespread following early unilateral enucleation, these localized changes highlight the importance of normal binocular vision for normal visual pathway development.

## Acknowledgements

This work was supported by NSERC Grants RPIN-05394 and RGPAS-477166 to BT.

## References

Ajina, S., Kennard, C., Rees, G., & Bridge, H. (2015). Motion area V5/MT+ response to global motion in the absence of V1 resembles early visual cortex. Brain, 138 (Pt 1), 164–178.

Alexander, A.L., Hurley, S.A., Samsonov, A.A., Adluru, N., Hosseinbor, A.P., Mossahebi, P., Tromp do, P.M., Zakszewski, E., & Field, A.S. (2011). Characterization of cerebral white matter properties using quantitative magnetic resonance imaging stains. Brain Connect, 1 (6), 423–446.

Allen, B., Schmitt, M.A., Kushner, B.J., & Rokers, B. (2018). Retinothalamic White Matter Abnormalities in Amblyopia. Invest Ophthalmol Vis Sci, 59 (2), 921–929.

Allen, B., Spiegel, D.P., Thompson, B., Pestilli, F., & Rokers, B. (2015). Altered white matter in early visual pathways of humans with amblyopia. Vision Res, 114, 48–55.

Barb, S.M., Rodriguez-Galindo, C., Wilson, M.W., Phillips, N.S., Zou, P., Scoggins, M.A., Li, Y., Qaddoumi, I., Helton, K.J., Bikhazi, G., Haik, B.G., & Ogg, R.J. (2011). Functional neuroimaging to characterize visual system development in children with retinoblastoma. Invest Ophthalmol Vis Sci, 52 (5), 2619–2626.

Behrens, T.E., Berg, H.J., Jbabdi, S., Rushworth, M.F., & Woolrich, M.W. (2007). Probabilistic diffusion tractography with multiple fibre orientations: What can we gain? Neuroimage, 34 (1), 144–155.

Behrens, T.E., Woolrich, M.W., Jenkinson, M., Johansen-Berg, H., Nunes, R.G., Clare, S., Matthews, P.M., Brady, J.M., & Smith, S.M. (2003). Characterization and propagation of uncertainty in diffusion-weighted MR imaging. Magn Reson Med, 50 (5), 1077–1088.

Born, R.T., & Bradley, D.C. (2005). Structure and function of visual area MT. Annu Rev Neurosci, 28, 157–189.

Desikan, R.S., Segonne, F., Fischl, B., Quinn, B.T., Dickerson, B.C., Blacker, D., Buckner, R.L., Dale, A.M., Maguire, R.P., Hyman, B.T., Albert, M.S., & Killiany, R.J. (2006). An automated labeling system for subdividing the human cerebral cortex on MRI scans into gyral based regions of interest. Neuroimage, 31 (3), 968–980.

Ding, K., Liu, Y., Yan, X., Lin, X., & Jiang, T. (2013). Altered functional connectivity of the primary visual cortex in subjects with amblyopia. Neural Plast, 2013, 612086.

Duan, Y., Norcia, A.M., Yeatman, J.D., & Mezer, A. (2015). The Structural Properties of Major White Matter Tracts in Strabismic Amblyopia. Invest Ophthalmol Vis Sci, 56 (9), 5152–5160.

Eickhoff, S.B., Heim, S., Zilles, K., & Amunts, K. (2006). Testing anatomically specified hypotheses in functional imaging using cytoarchitectonic maps. Neuroimage, 32 (2), 570–582.

Eickhoff, S.B., Paus, T., Caspers, S., Grosbras, M.H., Evans, A.C., Zilles, K., & Amunts, K. (2007). Assignment of functional activations to probabilistic cytoarchitectonic areas revisited. Neuroimage, 36 (3), 511–521.

Eickhoff, S.B., Stephan, K.E., Mohlberg, H., Grefkes, C., Fink, G.R., Amunts, K., & Zilles, K. (2005). A new SPM toolbox for combining probabilistic cytoarchitectonic maps and functional imaging data. Neuroimage, 25 (4), 1325–1335.

El-Shamayleh, Y., Kiorpes, L., Kohn, A., & Movshon, J.A. (2010). Visual motion processing by neurons in area MT of macaque monkeys with experimental amblyopia. J Neurosci, 30 (36), 12198–12209.

Frazier, J.A., Chiu, S., Breeze, J.L., Makris, N., Lange, N., Kennedy, D.N., Herbert, M.R., Bent, E.K., Koneru, V.K., Dieterich, M.E., Hodge, S.M., Rauch, S.L., Grant, P.E., Cohen, B.M., Seidman, L.J., Caviness, V.S., & Biederman, J. (2005). Structural brain magnetic resonance imaging of limbic and thalamic volumes in pediatric bipolar disorder. Am J Psychiatry, 162 (7), 1256–1265.

Gonzalez, E.G., Lillakas, L., Greenwald, N., Gallie, B.L., & Steinbach, M.J. (2014). Unaffected smooth pursuit but impaired motion perception in monocularly enucleated observers. Vision Res, 101, 151–157.

Ho, C.S., Giaschi, D.E., Boden, C., Dougherty, R., Cline, R., & Lyons, C. (2005). Deficient motion perception in the fellow eye of amblyopic children. Vision Res, 45 (12), 1615–1627.

Holmes, J.M., & Clarke, M.P. (2006). Amblyopia. Lancet, 367 (9519), 1343–1351.

Hoover, A.E., Harris, L.R., & Steeves, J.K. (2012). Sensory compensation in sound localization in people with one eye. Exp Brain Res, 216 (4), 565–574.

Jbabdi, S., Sotiropoulos, S.N., Savio, A.M., Grana, M., & Behrens, T.E. (2012). Model-based analysis of multishell diffusion MR data for tractography: how to get over fitting problems. Magn Reson Med, 68 (6), 1846–1855.

Jellison, B.J., Field, A.S., Medow, J., Lazar, M., Salamat, M.S., & Alexander, A.L. (2004). Diffusion tensor imaging of cerebral white matter: a pictorial review of physics, fiber tract anatomy, and tumor imaging patterns. AJNR Am J Neuroradiol, 25 (3), 356–369.

Jenkinson, M., Bannister, P., Brady, M., & Smith, S. (2002). Improved optimization for the robust and accurate linear registration and motion correction of brain images. Neuroimage, 17 (2), 825–841.

Jenkinson, M., Beckmann, C.F., Behrens, T.E., Woolrich, M.W., & Smith, S.M. (2012). Fsl. Neuroimage, 62 (2), 782–790.

Kelly, K.R., DeSimone, K.D., Gallie, B.L., & Steeves, J.K. (2015). Increased cortical surface area and gyrification following long-term survival from early monocular enucleation. Neuroimage Clin, 7, 297–305.

Kelly, K.R., Gallie, B.L., & Steeves, J.K. (2012). Impaired face processing in early monocular deprivation from enucleation. Optom Vis Sci, 89 (2), 137–147.

Kelly, K.R., Gallie, B.L., & Steeves, J.K.E. (2018). Early monocular enucleation selectively disrupts neural development of face perception in the occipital face area. Exp Eye Res,

Kelly, K.R., McKetton, L., Schneider, K.A., Gallie, B.L., & Steeves, J.K. (2014). Altered anterior visual system development following early monocular enucleation. Neuroimage Clin, 4, 72–81.

Kelly, K.R., Moro, S.S., & Steeves, J.K.E. (2012). Living with one eye: Plasticity in visual and auditory systems. In: J.K.E. Steeves, & L.R. Harris (Eds.), Plasticity in Sensory Systems (pp. 94–108). Cambridge, UK: Cambridge University Press.

Kelly, K.R., Zohar, S.R., Gallie, B.L., & Steeves, J.K. (2013). Impaired speed perception but intact luminance contrast perception in people with one eye. Invest Ophthalmol Vis Sci, 54 (4), 3058–3064.

Knyazeva, M.G. (2013). Splenium of corpus callosum: patterns of interhemispheric interaction in children and adults. Neural Plast, 2013, 639430.

Li, Q., Jiang, Q., Guo, M., Li, Q., Cai, C., & Yin, X. (2013). Grey and white matter changes in children with monocular amblyopia: voxel-based morphometry and diffusion tensor imaging study. Br J Ophthalmol, 97 (4), 524–529.

Li, X., Mullen, K.T., Thompson, B., & Hess, R.F. (2011). Effective connectivity anomalies in human amblyopia. Neuroimage, 54 (1), 505–516.

Liang, M., Xie, B., Yang, H., Yin, X., Wang, H., Yu, L., He, S., & Wang, J. (2017). Altered interhemispheric functional connectivity in patients with anisometropic and strabismic amblyopia: a resting-state fMRI study. Neuroradiology, 59 (5), 517–524.

Maurer, D., & Mc, K.S. (2018). Classification and diversity of amblyopia. Vis Neurosci, 35, E012.

Mendola, J.D., Conner, I.P., Roy, A., Chan, S.T., Schwartz, T.L., Odom, J.V., & Kwong, K.K. (2005). Voxel-based analysis of MRI detects abnormal visual cortex in children and adults with amblyopia. Hum Brain Mapp, 25 (2), 222–236.

Mendola, J.D., Lam, J., Rosenstein, M., Lewis, L.B., & Shmuel, A. (2018). Partial correlation analysis reveals abnormal retinotopically organized functional connectivity of visual areas in amblyopia. Neuroimage Clin, 18, 192–201.

Mitchell, D., & Sengpiel, F. (2018). Animal models of amblyopia. Vis Neurosci, 35, E017.

Moro, S.S., Kelly, K.R., McKetton, L., Gallie, B.L., & Steeves, J.K. (2015). Evidence of multisensory plasticity: Asymmetrical medial geniculate body in people with one eye. Neuroimage Clin, 9, 513–518.

Moro, S.S., & Steeves, J.K. (2012). No Colavita effect: equal auditory and visual processing in people with one eye. Exp Brain Res, 216 (3), 367–373.

Moro, S.S., & Steeves, J.K. (2013). No Colavita effect: increasing temporal load maintains equal auditory and visual processing in people with one eye. Neurosci Lett, 556, 186–190.

Mount, C.W., & Monje, M. (2017). Wrapped to Adapt: Experience-Dependent Myelination. Neuron, 95 (4), 743–756.

Nicholas, J.J., Heywood, C.A., & Cowey, A. (1996). Contrast sensitivity in one-eyed subjects. Vision Res, 36 (1), 175–180.

Nys, J., Scheyltjens, I., & Arckens, L. (2015). Visual system plasticity in mammals: the story of monocular enucleation-induced vision loss. Front Syst Neurosci, 9, 60.

Peiker, C., Wunderle, T., Eriksson, D., Schmidt, A., & Schmidt, K.E. (2013). An updated midline rule: visual callosal connections anticipate shape and motion in ongoing activity across the hemispheres. J Neurosci, 33 (46), 18036–18046.

Qi, S., Mu, Y.F., Cui, L.B., Li, R., Shi, M., Liu, Y., Xu, J.Q., Zhang, J., Yang, J., & Yin, H. (2016). Association of Optic Radiation Integrity with Cortical Thickness in Children with Anisometropic Amblyopia. Neurosci Bull, 32 (1), 51–60.

Restani, L., & Caleo, M. (2016). Reorganization of Visual Callosal Connections Following Alterations of Retinal Input and Brain Damage. Front Syst Neurosci, 10, 86.

Smith, S.M., Johansen-Berg, H., Jenkinson, M., Rueckert, D., Nichols, T.E., Miller, K.L., Robson, M.D., Jones, D.K., Klein, J.C., Bartsch, A.J., & Behrens, T.E. (2007). Acquisition and voxelwise analysis of multi-subject diffusion data with tract-based spatial statistics. Nat Protoc, 2 (3), 499–503.

Song, C., & Rees, G. (2018). Intra-hemispheric integration underlies perception of tilt illusion. Neuroimage, 175, 80–90.

Song, S.K., Sun, S.W., Ramsbottom, M.J., Chang, C., Russell, J., & Cross, A.H. (2002). Dysmyelination revealed through MRI as increased radial (but unchanged axial) diffusion of water. Neuroimage, 17 (3), 1429–1436.

Steeves, J.K., Gonzalez, E.G., Gallie, B.L., & Steinbach, M.J. (2002). Early unilateral enucleation disrupts motion processing. Vision Res, 42 (1), 143–150.

Steeves, J.K., Gonzalez, E.G., & Steinbach, M.J. (2008). Vision with one eye: a review of visual function following unilateral enucleation. Spat Vis, 21 (6), 509–529.

Thompson, B., Villeneuve, M.Y., Casanova, C., & Hess, R.F. (2012). Abnormal cortical processing of pattern motion in amblyopia: evidence from fMRI. Neuroimage, 60 (2), 1307–1315.

Wang, T., Li, Q., Guo, M., Peng, Y., Li, Q., Qin, W., & Yu, C. (2014). Abnormal functional connectivity density in children with anisometropic amblyopia at resting-state. Brain Res, 1563, 41–51.

Winkler, A.M., Ridgway, G.R., Webster, M.A., Smith, S.M., & Nichols, T.E. (2014). Permutation inference for the general linear model. Neuroimage, 92, 381–397.

Wong, N.A., Rafique, S.A., Kelly, K.R., Moro, S.S., Gallie, B.L., & Steeves, J.K.E. (2018). Altered white matter structure in the visual system following early monocular enucleation. Hum Brain Mapp, 39 (1), 133–144.

Wu, Y.C., Field, A.S., Duncan, I.D., Samsonov, A.A., Kondo, Y., Tudorascu, D., & Alexander, A.L. (2011). High b-value and diffusion tensor imaging in a canine model of dysmyelination and brain maturation. Neuroimage, 58 (3), 829–837.

Xiao, J.X., Xie, S., Ye, J.T., Liu, H.H., Gan, X.L., Gong, G.L., & Jiang, X.X. (2007). Detection of abnormal visual cortex in children with amblyopia by voxel-based morphometry. Am J Ophthalmol, 143 (3), 489–493.

Xie, S., Gong, G.L., Xiao, J.X., Ye, J.T., Liu, H.H., Gan, X.L., Jiang, Z.T., & Jiang, X.X. (2007). Underdevelopment of optic radiation in children with amblyopia: a tractography study. Am J Ophthalmol, 143 (4), 642–646.

Zhai, J., Chen, M., Liu, L., Zhao, X., Zhang, H., Luo, X., & Gao, J. (2013). Perceptual learning treatment in patients with anisometropic amblyopia: a neuroimaging study. Br J Ophthalmol, 97 (11), 1420–1424.

